# A recurrent circuit links antagonistic cerebellar modules during associative motor learning

**DOI:** 10.1101/2021.11.16.468438

**Authors:** Shogo Ohmae, Keiko Ohmae, Shane Heiney, Divya Subramanian, Javier Medina

## Abstract

The neural architecture of the cerebellum is thought to be specialized for performing supervised learning: specific error-related climbing fiber inputs are used to teach sensorimotor associations to small ensembles of Purkinje cells located in functionally distinct modules that operate independently of each other in a purely feedforward manner. Here, we test whether the basic operation of the cerebellum complies with this basic architecture in mice that learned a simple sensorimotor association during eyeblink conditioning. By recording Purkinje cells in different modules and testing whether their responses rely on recurrent circuits, our results reveal three operational principles about the functional organization of the cerebellum that stand in stark contrast to the conventional view: (1) Antagonistic organization, (2) Recurrent network dynamics, and (3) Intermodular communication. We propose that the neural architecture of the cerebellum implements these three operational principles to achieve optimal performance and solve a number of problems in motor control.

## INTRODUCTION

Recurrent connections are ubiquitous and feature prominently in the brain and in artificial neural networks, where they play a key role in a wide range of functions (Lisberger and Sejnowski, 1992; Douglas et al., 1995; Olshausen and Field, 1996; Hupe et al., 1998; Nakazawa et al., 2002; Wang, 2002; Porrill et al., 2004b; Mante et al., 2013; Bolding and Franks, 2018; Raymond and Medina, 2018; Wang et al., 2018; Peron et al., 2020; Kandel et al., 2021). These recurrent connections often have a closed-loop or reciprocal organization, in which the final output of one functional module is fed back to the input layer of the same module (closed LOOP architecture), thus ensuring that signals are kept private and processed independently in anatomically segregated channels (Douglas et al., 1995; Olshausen and Field, 1996; Hupe et al., 1998; Nakazawa et al., 2002; Mante et al., 2013; Wang et al., 2018). However, recurrent connections can also provide a substrate for coupling the function of different modules together, by linking the output of one module to the input of another (open SPIRAL architecture) (Haber et al., 2000; Haruno and Kawato, 2006). Defining the role of these two types of recurrent circuits is a crucial step for understanding the global operation of neural systems in many parts of the brain. In this paper, we examine how recurrent connectivity impacts signal processing in the cerebellum, a brain system that incorporates both LOOP and SPIRAL circuits in its underlying neural architecture.

The microcircuit of the cerebellum is well known for its critical role in adaptive motor control, and for its highly stereotyped connectivity and modular organization (Eccles et al., 1967; Apps and Garwicz, 2005; Apps et al., 2018). Each individual cerebellar module contains a group of sagittally oriented Purkinje cells that receive climbing fiber inputs with the same receptive field, and that in turn project to neurons located in discrete and compartmentalized areas of the deep cerebellar nuclei (DCN). Previous analyses of motor-related responses in the cerebellum have helped define how signals are processed along the feedforward stream that links Purkinje cells to DCN neurons (Raymond and Medina, 2018). In contrast, there is a complete lack of information about the contribution of feedback signals flowing in the reverse direction, even though anatomical studies have revealed LOOP and SPIRAL circuits connecting DCN and Purkinje cells (Apps and Garwicz, 2005; Houck and Person, 2014; Ankri et al., 2015; Houck and Person, 2015; Gao et al., 2016), and recent findings indicate that these recurrent pathways are functional and may play an important role in gain control and pattern completion of learned movements (Gao et al., 2016; Khilkevich et al., 2018a; Khilkevich et al., 2018b).

Our experiments provide a first look at the role of recurrent circuits in generating motor-related signals in Purkinje cells. We trained mice on a Pavlovian eyeblink conditioning task and analyzed how the activity of Purkinje cells was altered during brief optogenetic inhibition of recurrent pathways originating in the rostral anterior interpositus nucleus (rAIN), a subregion of the DCN that is essential for performance of the conditioned eyelid movement (conditioned response, CR) (Heiney et al., 2014b). To tease apart the relative contributions of LOOP and SPIRAL circuits, we measured the impact of rAIN photoinhibition on the activity of rAIN-projecting Purkinje cells responsible for CR generation (Center Purkinje cells, CTR-PCs), as well as Purkinje cells in neighboring cerebellar modules without projections to the rAIN region (Surround Purkinje cells, SND-PCs).

## RESULTS

In all the experiments described below, we recorded the extracellular spiking activity of Purkinje cells in mice that had been thoroughly trained on a Pavlovian eyeblink conditioning task. At the time of the Purkinje cell recordings, all the mice (n=19) had learned to make a blink (conditioned response, CR) in response to a warning cue (conditioned stimulus, CS) that was repeatedly paired with an aversive airpuff directed at the eye (unconditioned stimulus, US; see Online Methods for details).

### CR-related responses of Purkinje cells in CENTER module

We recorded the activity of Purkinje cells located in specific modules of the cerebellar cortex, which could be distinguished from each other based on the receptive field properties of their climbing fiber input (Apps and Garwicz, 2005; Mostofi et al., 2010; Heiney et al., 2014a). Purkinje cells in the CENTER module (CTR-PCs) fired a climbing fiber-driven complex spike (CSpk) with a short-latency after delivering an unexpected airpuff to the mouse’s eye (Fig. 1a,e; Supplementary Fig. 1a for individual cells; CSpk latency: 11 to 42 milliseconds). The eyepuff also produced a modulation of simple spikes (SS) in CTR-PCs. This SS response to the eyepuff often included an initial inhibition (Fig. 1a,g; Supplementary Fig. 1b), which may be driven by activation of inputs from molecular layer interneurons (Ekerot and Jorntell, 2001), as well as by the well-documented suppression of SS activity that follows the CSpk (Eccles et al., 1966; Sato et al., 1992)(Fig. 1a, inset).

**Figure 1.**
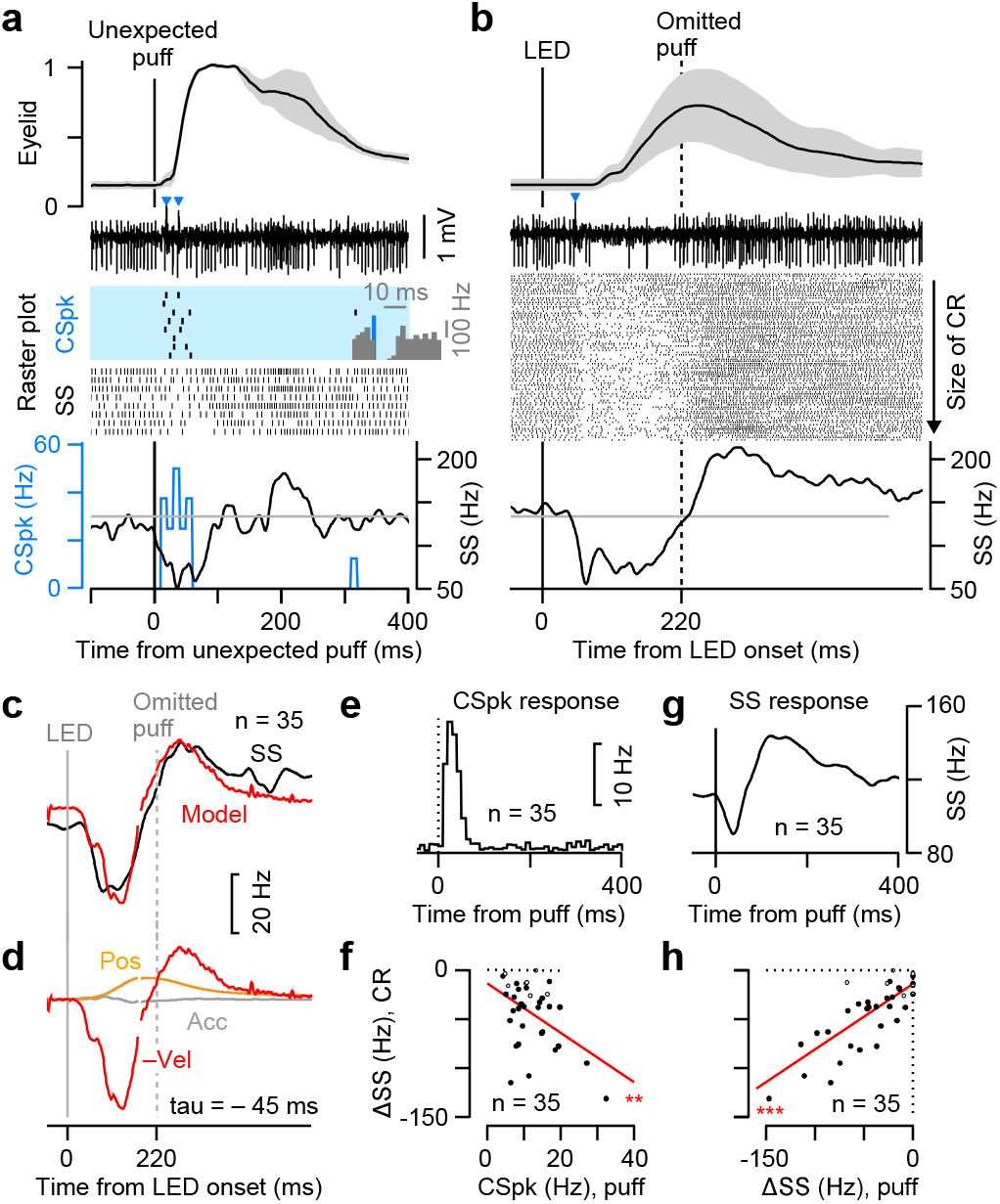
CR-related activity of CENTER Purkinje cells. **a-b**, Eyelid movements (mean (line) ± s.d. (shaded region)), raw extracellularly recorded voltage signal in an example trial, and detected SSs (black) and CSpks (blue; histogram of SS suppression after CSpks in the inset) fired by a representative CENTER Purkinje cell in trials with unexpected delivery of eyepuff (**a**) and in eyeblink conditioning trials with LED CS without eyepuff US (**b**). **c**, Model prediction for population averaged SS firing of CENTER Purkinje cells with the kinematics of the eyelid movement: SS(t) = K1* position(t-tau) + K2 * velocity(t-tau) + K3 * acceleration (t-tau) + K4. **d**, Contribution of each kinematic component to the predictive model. **e,g**, Population response of CSpks (**e**) and SSs (**g**) of CENTER Purkinje cells to unexpected eyepuffs. **f,h**, Maximum depth of rate change of SS firing during CR performance relative to the Cspk probability (**f**) and to the suppression of SS firing (**h**) after unexpected eyepuff, plotted for all individual Purkinje cells (Pearson’s correlation coefficients; **f**, n = 35 cells, r = 0.47, P = 0.0045; **h**, n = 35 cells, r = 0.81, P < 0.001). SS suppression during CR was significant in 28 cells (closed dots; Wilcoxon signed rank test) but not in 7 cells (open dots). ** P < 0.01, *** P < 0.001.

We examined the CR-related activity of CTR-PCs after thoroughly training the mice in a Pavlovian eyeblink conditioning task in which a cue stimulus (an LED pulse) was repeatedly paired with an eyepuff at a specific interval (interstimulus interval, ISI). To reveal the entire time course of CR-related activity without contamination from reflex blinks, we limited our analysis to test trials in which the cue stimulus was presented by itself without the eyepuff. As in previous experiments (Hesslow and Ivarsson, 1994; Jirenhed and Hesslow, 2011; Halverson et al., 2015; ten Brinke et al., 2015), the most common response we observed in CTR-PCs was a reduction of SS firing that persisted until the time that the eyepuff would have been presented (Fig. 1b-c; Supplementary Fig. 1c), regardless of the ISI used during training (Supplementary Fig. 2a,c). The time course of the SS firing rate modulation matched the inverted velocity profile of the CR (Fig. 1c,d; Supplementary Fig. 2a,c), and in many cells (n = 21/35) the depth of the modulation was proportional to the speed of the CR on individual trials (Supplementary Fig 3a-c). Taking into account that the average SS activity of the population (n = 35) preceded the eyelid kinematics (tau = -45 ms; Fig. 1d) and given that Purkinje cells are inhibitory (Ito et al., 1964; Eccles et al., 1967), our results are consistent with previous studies suggesting that the reduction in the SS activity of CTR-PCs is partly responsible for driving the CR (Perrett et al., 1993; Gerwig et al., 2005; Mostofi et al., 2010; Steinmetz and Freeman, 2014), via disinhibition of downstream neurons in the cerebellar nucleus (Heiney et al., 2014a).

Although the majority of CTR-PCs reduced their SS firing rates during the CR (n = 28/35), the extent of the reduction varied considerably from cell to cell (Fig. 1f,h; Supplementary Fig. 1c). Further analyses revealed that the depth of the CR-related reduction in SS activity was strongly correlated with two features about the response of each cell to the unexpected presentations of the eyepuff stimulus: CTR-PCs with the largest reductions in CR-related SS activity were the ones that responded to the eyepuff with a highly reliable Cspk (Fig. 1f; Supplementary Fig. 1a) and a large suppression of SS activity (Fig. 1h; Supplementary Fig. 1b). Because the eyepuff serves as the instructive stimulus during eyeblink conditioning, these results are consistent with theories that assign a teaching role to either CSpk’s (Marr, 1969; Raymond et al., 1996; De Zeeuw et al., 1998; Doya, 2000; Ito, 2006; Dean et al., 2010; Simpson et al., 2011) or SS’s (Miles and Lisberger, 1981; Raymond and Lisberger, 1998; Medina and Mauk, 1999; Medina, 2011; Popa et al., 2015).

### Contribution of recurrent LOOP circuits to CR-related responses of CTR-PCs

Previous studies have demonstrated that similar to what we found for CTR-PCs, the SS activity of Purkinje cells is often correlated with the kinematic parameters of future movements (Shidara et al., 1993; Medina and Lisberger, 2009; Herzfeld et al., 2015; Chen et al., 2016; Hong et al., 2016; Dugue et al., 2017; Sun et al., 2017; Popa et al., 2019). However, the relative contribution of different input streams to this motor-related response has not been investigated and remains a matter of considerable debate (Medina, 2011; Houck and Person, 2014; Raymond and Medina, 2018). We used an optogenetic approach to examine whether recurrent LOOP activity was necessary for the CR-related SS response of CTR-PCs (Fig. 2a). To silence recurrent LOOP pathways originating in the CENTER module, we photostimulated inhibitory Purkinje cell axon terminals of PCP2-ChR2 mice via an optical fiber placed in the rostrolateral anterior interpositus nucleus (rAIN, Fig. 2b; Supplementary Fig. 4a), which is the region of the cerebellar nuclei (CTR-CN) that drives the CR (McCormick and Thompson, 1984; Yeo et al., 1985; Heiney et al., 2014a; Heiney et al., 2014b) and is thought to receive Purkinje cell projections from areas of cerebellar cortex where CTR-PCs were located (Apps and Hawkes, 2009; Mostofi et al., 2010; Heiney et al., 2014a; Ten Brinke et al., 2017; Heiney et al., 2021)(Supplementary Fig. 5). In every optogenetic experiment, we performed two tests to confirm the correct placement of the optical fiber, and data was analyzed only if we could verify that: (1) photostimulation suppressed CRs (Supplementary Fig. 4b,c), as expected during successful inhibition of rAIN neurons in the CENTER module (Heiney et al., 2014b), and (2) photostimulation resulted in antidromic activation of the CTR-PC being recorded (Fig. 2c; Supplementary Fig. 4d,e), as expected for Purkinje cells belonging to the same module as the neurons of the cerebellar nuclei where the optical fiber was located.

**Figure 2.**
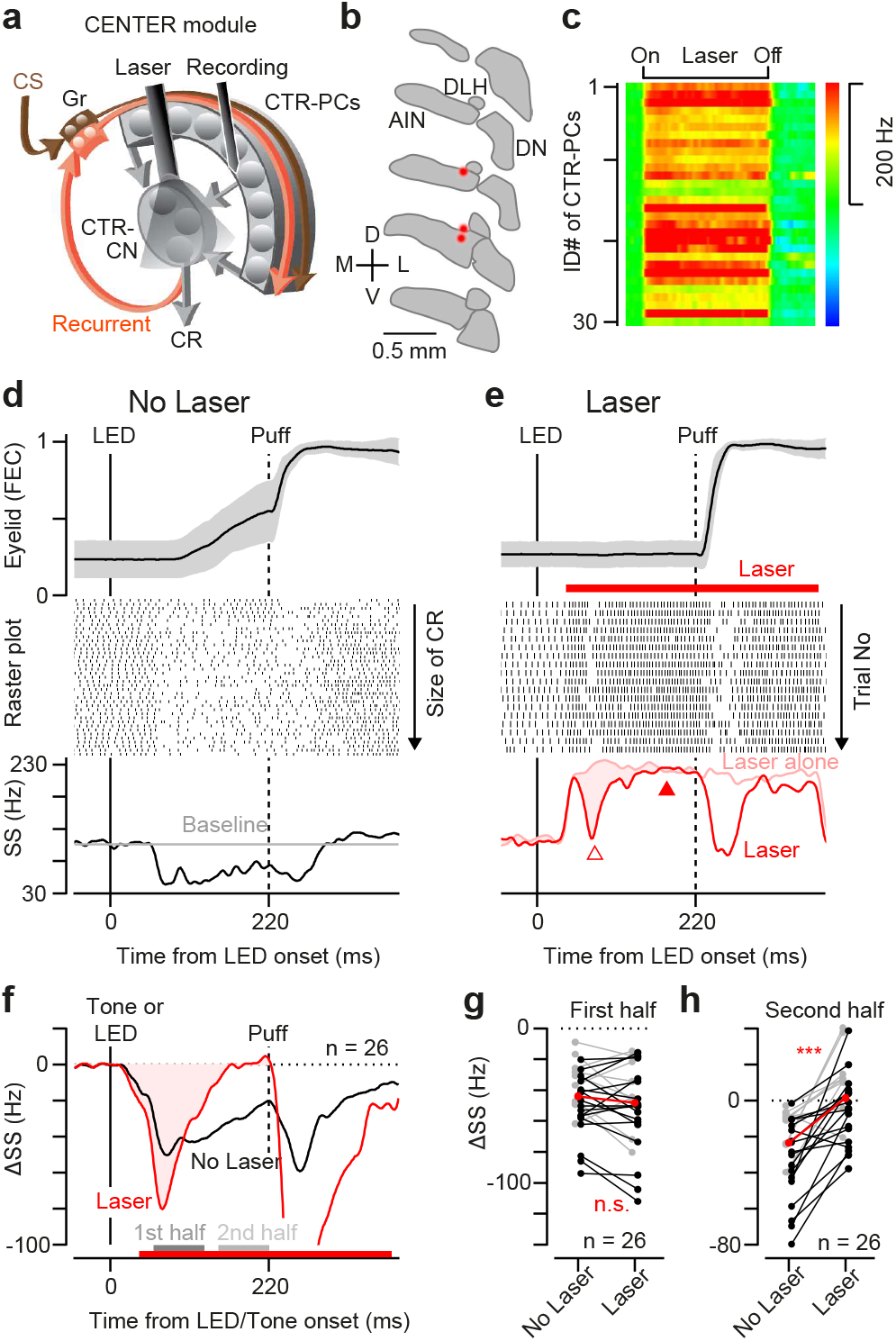
Effect of rAIN photo-inhibition on CENTER Purkinje cells. **a**, Experimental design for recording CENTER Purkinje cells during rAIN inhibition by GABAergic-terminal photostimulation. **b**, Locations of optic fiber tips (red dots; n = 3/4 mice) on the contours of coronal sections of the cerebellar nuclei (every 80 μm from rostral to caudal). AIN, anterior interpositus nucleus; DN, dentate nucleus; DLH, dorsolateral hump. **c**, Antidromic response of CENTER Purkinje cells during photostimulation of Purkinje cell axon terminals in rAIN. **d,e**, Eyelid movements (mean ± s.d. (shaded region)) and simultaneously recorded SS responses to LED CS of a representative CENTER Purkinje cell in the absence (**d**) and presence (**e**) of the GABAergic terminal photostimulation. The CR-related firing modulation during rAIN photoinhibition (red shaded area) was calculated by subtracting the response in trials with ‘laser alone’ from the response in conditioning trials with ‘laser’. **f**, Population-averaged CR-related SS activity in the absence (black) and presence (red) of rAIN photoinhibition. **g,h**, Comparison of firing modulation in trials with and without laser for individual Purkinje cells and the median (red) after the LED CS (black) and TONE CS (gray) during the first half (**g**, dark gray in **f**) and the second half (**h**, light gray in **f**) of the CR period (Wilcoxon signed rank test; **g**, n = 26 cells, P = 0.18; **h**, n = 26 cells, P < 0.001). n.s. not significant, *** P < 0.001.

We recorded the activity of CTR-PCs (n = 18) during conditioning sessions in which we alternated normal trials (Fig. 2d) and trials with laser-driven photoinhibition of the rAIN region during the CR period (Fig. 2e; laser duration 350 ms, onset 40 ms after LED CS onset or 10 ms after tone CS onset). Despite the antidromic activation resulting from the stimulation of Purkinje cell axons in the laser trials (Fig. 2c,e), a reduction in SS activity was clearly visible following presentation of the LED stimulus (Fig. 2e-f, ‘Laser’), even after discarding all trials in which there was a Cspk in the CR period (Supplementary Fig. 6, ‘Laser Cspk-‘). Analysis of the time course of this reduction in SS firing, which was calculated by subtracting the SS response in conditioning trials with laser from the SS response to the laser alone (Fig. 2e, shaded area), revealed that the initial component was similar to what we observed in normal conditioning trials without laser stimulation (Fig. 2f,g). However, the reduction of SS firing did not last as long and terminated significantly earlier in laser trials (Fig. 2f,h). The same results were obtained when we used a tone instead of the LED as the CS (Fig. 2g,h, gray circles). This finding suggests that the SS response of CTR-PCs during the CR can be divided into two components: an initial reduction in SS firing that is brief and driven by activation of feedforward signals related to the cue stimulus (Fig. 2a, ‘CS’), followed by a more prolonged SS suppression under the control of recurrent circuits originating in the rAIN region of the cerebellar nuclei (Fig. 2a, ‘Recurrent’).

### CR-related responses of Purkinje cells in SURROUND modules

In addition to CTR-PCs, we also recorded the activity of Purkinje cells in areas of the cerebellar cortex surrounding the CENTER module (SND-PCs, Supplementary Fig. 5 shows specific locations confirmed histologically for 5 example cells). SND-PCs were defined as Purkinje cells that were recorded within a 500 µm radius of CTR-PCs but lacked a Cspk response to the eyepuff stimulus (Fig. 3a,e, Supplementary Fig. 1a), indicating that their climbing fiber receptive field was located outside the periocular region. To verify that our electrophysiological recordings of SND-PCs belonged to Purkinje cells and not to one of the other cell types in the cerebellar cortex, we confirmed that the cells fired occasional spontaneous Cspks at the normal low rate of ∼1 Hz during baseline periods (Fig. 3a,b,e), that each spontaneous Cspk was followed by the characteristic pause in SSs (Fig. 3a, inset), and that SS activity was strongly inhibited by optogenetic stimulation of nearby molecular layer interneurons (Supplementary Fig. 7).

**Figure 3.**
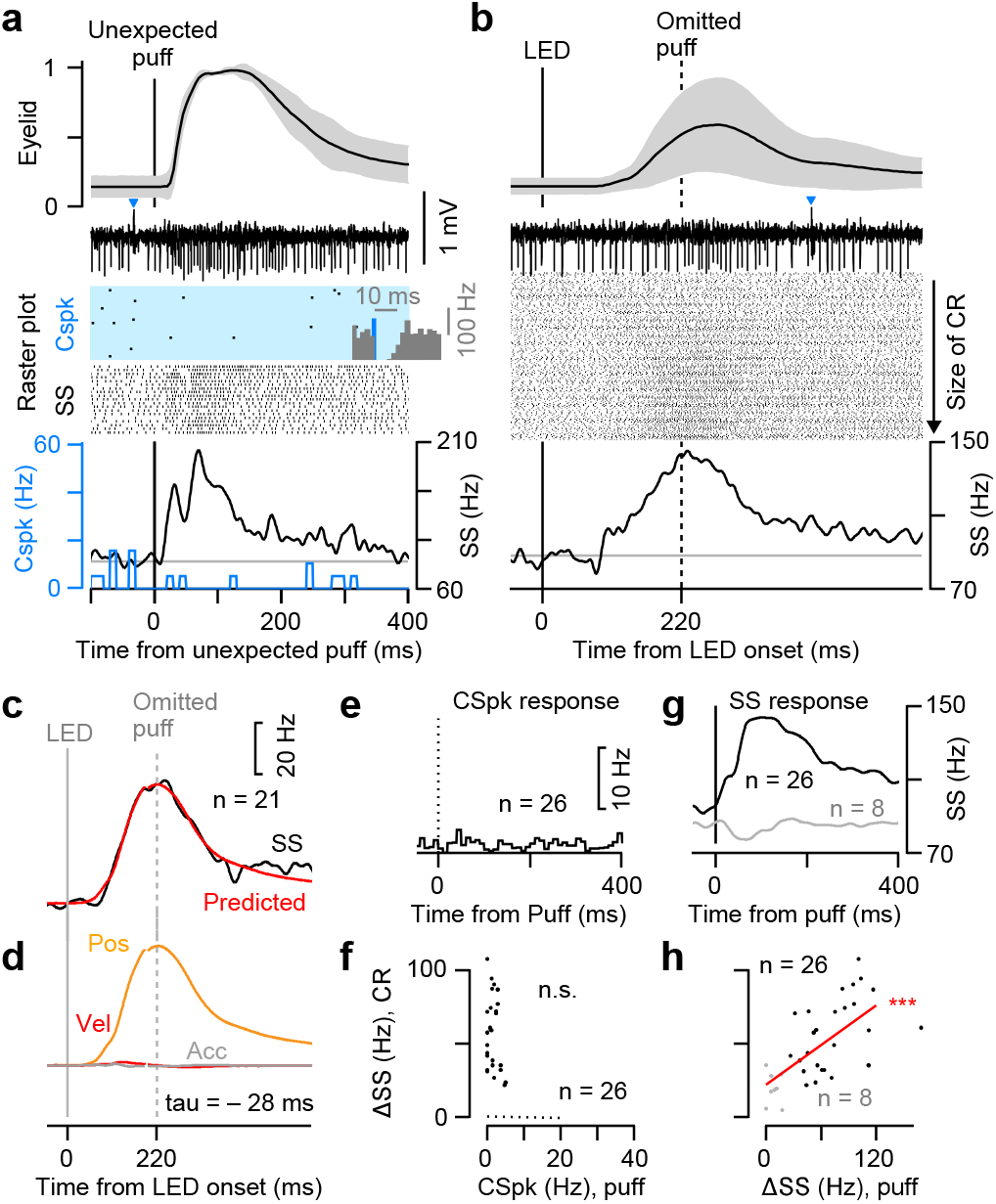
CR-related activity of SURROUND Purkinje cells. **a**-**b**, Eyelid movements (mean (line) ± s.d. (shaded region)), raw extracellularly recorded voltage signal in an example trial, and detected SSs (black) and CSpks (blue; histogram of SS suppression after CSpks in the inset) fired by a representative SURROUND Purkinje cell in trials with unexpected delivery of eyepuff (a) and in eyeblink conditioning trials with LED CS without eyepuff US (**b**). **c**, Model prediction for population averaged SS firing of SURROUND Purkinje cells with the kinematics of the eyelid movement: SS(t) = K1* position(t-tau) + K2 * velocity(t-tau) + K3 * acceleration (t-tau) + K4. **d**, Contribution of each kinematic component to the predictive model. **e,g**, Population response of CSpks (**e**) and SSs (**g**) of SURROUND Purkinje cells to unexpected eyepuffs. **f,h**, Maximum rise of SS firing rate during CR performance relative to the Cspk probability (**f**) and to the facilitation of SS firing (**h**) after unexpected eyepuff, plotted for all individual Purkinje cells (Pearson’s correlation coefficients, **f**, n = 26 cells, r = 0.25, P = 0.22; **h**, n = 34 cells, r = 0.67, P < 0.001). In panels **g,h**, data for nearby Purkinje cells without SS response to the unexpected eyepuff (n = 8) is shown in gray. n.s. not significant, *** P < 0.001.

We found a number of critical differences between the SS responses of SND-PCs and CTR-PCs: (1) The baseline SS firing rate was higher in SND-PCs than in CTR-PCs (Supplementary Fig. 8), (2) the SS firing rate of most SND-PCs increased following the unexpected eyepuff stimulus (26/34 cells, Fig. 3g), without displaying any signs of the short-latency inhibition that was characteristic of CTR-PCs (Supplementary Fig. 1b), (3) SS firing rate went up during the CR period (Fig. 3b,c,f,h), which is opposite to the downward modulation we observed in CTR-PCs (Supplementary Fig. 1c), (4) Whereas we had found a clear link between the size of the CR-related SS response of individual CTR-PCs and the size of both their SS and Cspk response to the eyepuff (Fig. 1f,h), only the relationship between the SS responses was preserved in SND-PCs (Fig. 3f,h), (5) The SS response of SND-PCs was strongly correlated with the eyelid position during the CR (Fig. 3d, Supplementary Fig. 3d-f), regardless of the ISI used in training (Supplementary Fig. 2b,d). This position-related modulation led the eyelid kinematics by ∼28 ms on average (Fig. 3d), which contrasts with the longer leading velocity-related modulation of the CTR-PC population (Fig. 1d) and results in SND-PCs reaching their maximum SS activation after the CR-related reduction in the SS firing rate of CTR-PCs has fully recovered. Overall, our results reveal an antagonistic population code for movement parameters in the cerebellar cortex, which is based on precisely coordinating the opposing SS responses of Purkinje cells in the CENTER and SURROUND modules.

### Contribution of recurrent SPIRAL circuits to CR-related responses of SND-PCs

Anatomical studies have hinted at the possibility that distinct regions of the cerebellar cortex could be functionally linked via SPIRAL circuits that directly or indirectly connect the output layer of one cerebellar module with the input layer of another (Tsukahara et al., 1983; Trott et al., 1998a, b; Ankri et al., 2015; Houck and Person, 2015; Gao et al., 2016). To formally test this hypothesis, we examined whether the output coming out of the CENTER module is necessary for the CR-related SS responses of Purkinje cells in SURROUND modules by recording the activity of SND-PCs while optogenetically inhibiting the rAIN region of the cerebellar nuclei (Fig. 4a). As in the experiments shown in Fig. 2, we confirmed that the optical fiber was correctly placed in rAIN by verifying that CRs were suppressed during laser stimulation (Supplementary Fig. 4b,c). Antidromic activation resulting from photostimulation of axons and terminals was only apparent in a few of the recorded SND-PCs (SS firing rate increased by >50% in only 2/14 cells), as may be expected for Purkinje cells that mainly project to areas of the cerebellar nuclei outside the rAIN region (Fig. 4b; Supplementary Fig. 4d,e).

**Figure 4.**
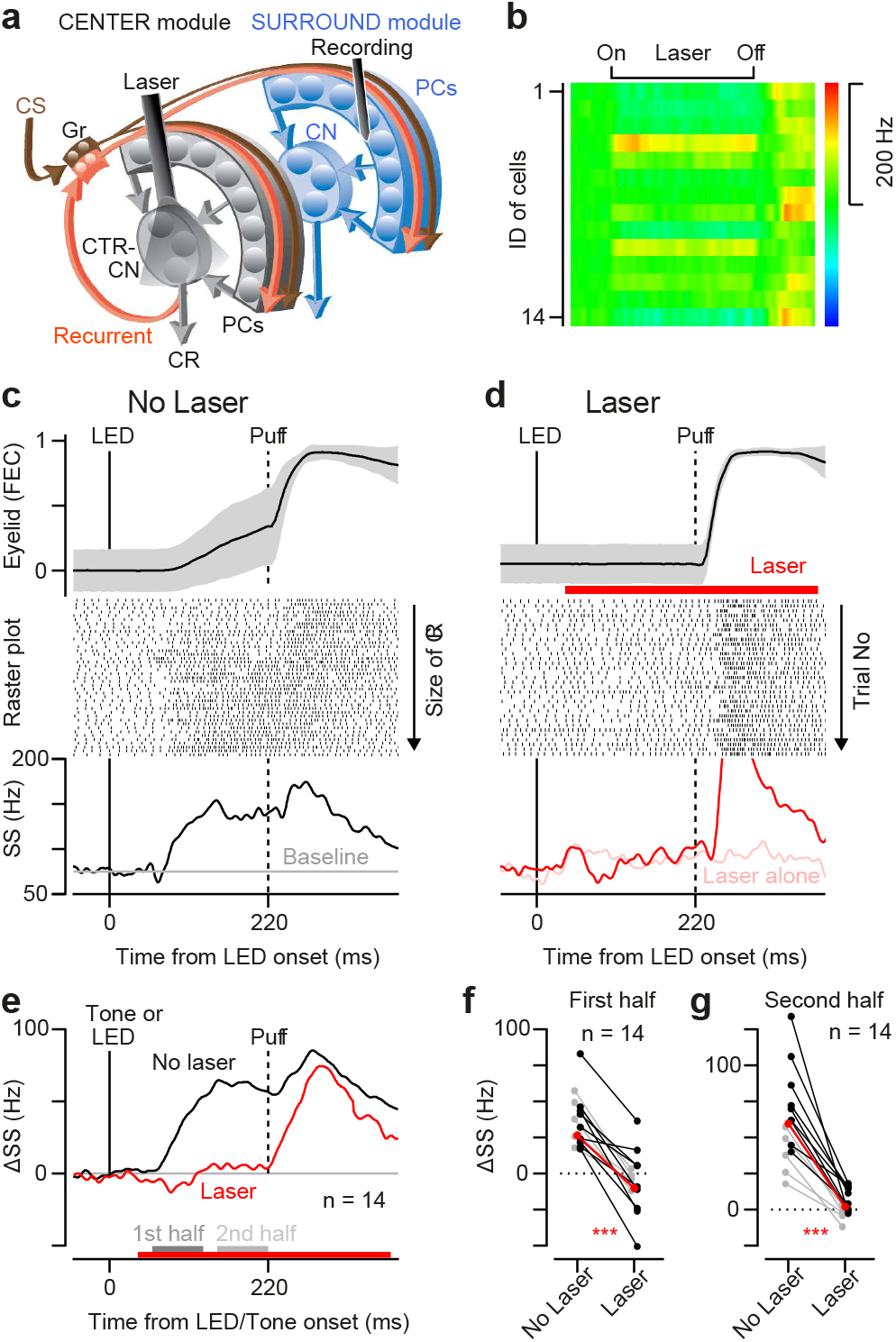
Effect of rAIN photo-inhibition on SURROUND Purkinje cells. **a**, Experimental design for recording SURROUND Purkinje cells during rAIN inhibition by GABAergic-terminal photostimulation. **b**, Antidromic response of SURROUND Purkinje cells during stimulation of Purkinje cell axon terminals in rAIN. **c,d**, Eyelid movements (mean ± s.d. (shaded region)) and simultaneously recorded SS responses to LED CS of a representative SURROUND Purkinje cell in the absence (**c**) and presence (**d**) of rAIN photoinhibition. **e**, Population-averaged modulation of SS firing after LED/TONE CS in the absence (black) and presence (red) of rAIN photoinhibition. **f,g**, Comparison of firing modulation in trials with and without laser for individual Purkinje cells and the median (red) after the LED CS (black) and TONE CS (gray) during the first half (**f**) and the second half (**g**) of the CR period (Wilcoxon signed rank test; **f**, n = 14 cells, P < 0.001; **g**, n = 14 cells, P < 0.001). *** P < 0.001.

We found that laser photostimulation completely eliminated the CR-related SS response of SND-PCs (Fig. 4c-e), including both the early (Fig. 4f) and late components (Fig. 4g). Two additional observations indicate that this effect was caused by a specific loss of CR-related SS activity and not by inhibition of SND-PCs via direct laser-driven activation of axon collaterals from nearby Purkinje cells (Witter et al., 2016), or by rendering SND-PCs generally unresponsive to all inputs: First, photostimulation had very little effect on the baseline spontaneous rate of SS activity of most SND-PCs (Fig. 4b,d-e; Supplementary Fig. 4d,e), and second, SND-PCs retained their normal SS response to the eyepuff stimulus during the photostimulation pulse (Fig. 4d,e; Supplementary Fig. 9). Thus, the movement-related SS activity that Purkinje cells in the SURROUND modules generate during performance of the CR is entirely driven by activation of direct or indirect recurrent SPIRAL circuits originating in rAIN, i.e. the output layer of the CENTER module.

### Flow of CR-related motor signals in the cerebellum

To gain further insight into the flow of signals and sequential recruitment of different nodes in the cerebellar network during motor performance, we compared the latency of the CR-related responses of CTR-PCs, SND-PCs and a previously recorded population of rAIN neurons (CTR-CNs) (Ten Brinke et al., 2017). When the activity of all cells was aligned on the time of the CR onset, it became clear that the population averaged response of CTR-PCs started first, followed by CTR-CNs in rAIN, and finally SND-PCs (Fig. 5a). The analysis of individual cells shown in Figure 5b,c (see Online Methods) confirmed that the onset of the response was significantly earlier in CTR-PCs (median latency = 36 ms before CR) than in CTR-CNs (median latency = 18 ms before CR) and SND-PCs (median latency = 4 ms before CR). The difference between the onset of the response in CTR-CNs and SND-PCs was small, however, and only marginally significant (Fig. 5b).

**Figure 5.**
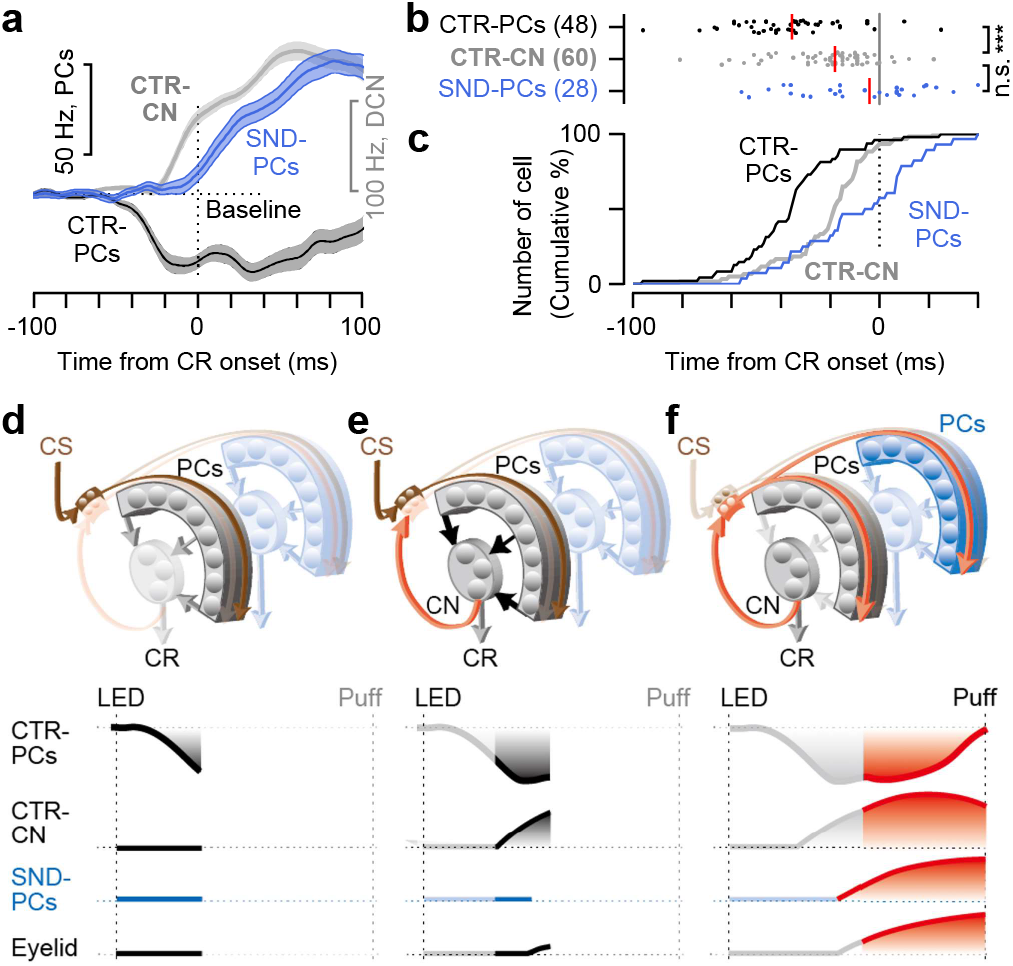
Sequential activation of CENTER and SURROUND modules. **a**, Population averaged firing rate of CTR-PCs, CTR-CN neurons, and SND-PCs, aligned to the CR onset. **b**, Onset times for the response of individual neurons (dots) and the median (red) (Kruskal-Wallis ANOVA, Chi-sq = 27.6, d.f. = 2, P < 0.001; Fisher’s least significant difference procedure multiple comparison, P < 0.001 between CTR-PCs and CTR-CN, P = 0.087 between CTR-CN and SND-PCs). **c**, Cumulative frequency histogram for the onset latency data in **b**. **d**, CS input (brown arrow) triggers the downward firing modulation in the CTR-PCs (black). **e**, CTR-PCs disinhibit CTR-CN neurons (black arrows), to drive the initiation of the CR. **f**, The CTR-CN provides recurrent LOOP and SPIRAL signal streams (red arrows) to maintain the down modulation of CTR-PCs and to drive the up modulation of SND-PCs for the remainder of the CR production.

Altogether, our findings suggest the following hypothesis about the dynamical evolution of movement-related signals in the cerebellum and their contribution to the responses of Purkinje cells in CENTER and SURROUND modules during performance of the CR (Fig. 5d-f): First, the CS signal is transmitted via granule cells to CTR-PCs, triggering a reduction in their SS firing rate that does not require activation of recurrent inputs (Fig. 5d). This reduction in CTR-PC activity causes disinhibition of rAIN neurons in the CENTER module (Heiney et al., 2014a), producing a signal that serves to initiate the CR (McCormick and Thompson, 1984; Yeo et al., 1985; Heiney et al., 2014a; Heiney et al., 2014b) and to activate recurrent LOOP and SPIRAL circuits projecting back to the cerebellar cortex (Fig. 5e). The recurrent signal reaches the cerebellar cortex and is distributed to CENTER and SURROUND modules very quickly and before the CR onset (<15 ms based on latency analysis in Fig. 5b and Supplementary Fig. 10), perhaps via direct projections from the cerebellar nuclei (Apps and Garwicz, 2005; Houck and Person, 2014, 2015; Gao et al., 2016). This activation of recurrent LOOP and SPIRAL circuits leads to a reverberation of signals in the cerebellum that is necessary for generating the motor-related responses of CTR-PCs and SND-PCs and for driving the CR (Fig. 5f).

## DISCUSSION

We have elucidated the contribution of recurrent circuits to the motor-related activity of Purkinje cells in two different modules of the cerebellar cortex: (1) Purkinje cells in the CENTER module (CTR-PCs) show a reduction in activity that is proportional to the velocity of a learned eyelid movement (conditioned response, CR). The latter part of this velocity-related response requires activation of recurrent LOOP pathways originating in the rostral anterior interpositus nucleus (rAIN), which is the area of the cerebellar nuclei (CN) where CTR-PCs project. (2) Purkinje cells in SURROUND modules (SND-PCs) show an increase in activity that is antagonistic to the response of CTR-PCs and is proportional to the position of the eyelid during the CR. This position-related response is entirely driven by recurrent SPIRAL pathways that link rAIN neurons of the CENTER module with SND-PCs of the SURROUND module. As discussed below, these findings suggest the need to make a number of substantial revisions to current theories about the functional organization of the cerebellum.

### Recurrent architecture

One classical view that should be revised based on our results is the idea that the cerebellum is a feed-forward network, i.e., an idea that is the foundation for computational theories of the cerebellum based on machine learning principles (Shadmehr, 2020) and the perceptron model (Eccles et al., 1967; Marr, 1969; Albus, 1971; Ito, 1972; Brunel et al., 2004). Despite plenty of anatomical evidence for the existence of feedback pathways from CN to PCs (disynaptic via granule cells, or trisynaptic additionally via Golgi cells or via precerebellar structures) (Apps and Garwicz, 2005; Ankri et al., 2015; Houck and Person, 2015; Raymond and Medina, 2018), few studies have examined its functional significance (Gao et al., 2016). Today, the classical view of the cerebellum as a feedforward network dominates, is introduced in textbooks (Kandel et al., 2021), and has even found its way into the mainstream media (Wikipedia). The classical view provides a foundation for discussing differences in functional specialization between the cerebellum and other brain regions with abundant recurrent connections, such as the cerebral cortex (e.g. in an Nature article (Koch, 2018)). In contrast to this classical view, we now demonstrate that recurrent signals from the CN play a key role in the signal processing of PCs. Our results also suggest that the recurrent signals are sent by fast feedback pathways, such as those that link the CN and the cerebellar cortex directly (Apps and Garwicz, 2005; Ankri et al., 2015; Houck and Person, 2015; Gao et al., 2016). Regardless of whether the recurrent signals are sent back to the cerebellar cortex from the CN directly or indirectly, our findings make it clear that feedforward processing, which is one of the principles of cerebellar computation, must be revised to include the extra operational power provided by a recurrent neural network architecture.

### Interdependent modules

Another classical view that should be revised in light of our results is the idea that the modules of the cerebellum are independent in the sense that the output of one module (i.e. CN signal) does not affect the computation of another module. This idea is the basis of a theory in which the cerebellum is functionally divided into several thousands of independent computational modules or units that provide massive parallel processing power (Apps and Garwicz, 2005; Apps et al., 2018). The massively parallel and independent modular architecture is another one of the features commonly brought up to highlight the differences in computational styles between the cerebellum and the cerebral cortex, in which strong horizontal connectivity is assumed to play a key role in allowing inter-modular interactions that support higher-level cognitive processing (Koch, 2018). In contrast to the view that cerebellar modules operate independently, there is recent evidence indicating that communication between different modules can be flexibly regulated to achieve optimal behavioral performance (Valera et al., 2016; Apps et al., 2018; Heiney et al., 2021). Our data reveals a strong inter-modular interaction in which recurrent SPIRAL pathways originating in the output layer of one module are responsible for driving the functional response of neurons in different modules. Therefore, we propose that the cerebellum should be viewed as a specific type of recurrent neural network, i.e., an interdependent-modular recurrent neural network with both LOOP and SPIRAL connections.

### Antagonistic CENTER/SURROUND organization

During cerebellar-dependent learning tasks, it is usual to find separate groups of PC populations with antagonistic sensorimotor-related responses (Jorntell and Ekerot, 2002; Medina and Lisberger, 2008; Herzfeld et al., 2015; Chabrol et al., 2019; Laurens and Angelaki, 2020). In vitro, an antagonistic center/surround organization was found to arise from direct excitatory input from granule cells in the center and indirect inhibitory input via molecular layer interneurons in the surround (Dizon and Khodakhah, 2011), while in awake behaving animals, microzones with antagonistic PCs have been defined based on the activity of their climbing fiber inputs during learning (Medina and Lisberger, 2008; Herzfeld et al., 2015, 2018; Chabrol et al., 2019). Our results demonstrate that antagonistic PC responses are also generated in CENTER and SURROUND modules during cerebellum-dependent eyeblink conditioning, lending further support to the idea that this antagonistic organization is one of the fundamental principles underlying cerebellar learning. A new theory has proposed that antagonistic zones in the cerebellar cortex (i.e. Downbound & Upbound zones) correspond to the zoning of PC subtypes classified by marker genes such as zebrin (Zhou et al., 2014; De Zeeuw, 2021). We did not find any evidence that CTR-PCs and SND-PCs belong to different zebrin bands. However, considering that the zebrin bands in the walls of the primary fissure are not as easily discernable as in other regions of the cerebellum (Sugihara and Quy, 2007; Fujita et al., 2014) and that zebrin is only one of many possible markers of PC subtypes (Sillitoe and Joyner, 2007; Apps and Hawkes, 2009), it would be premature to conclude that CTR-PCs and SND-PCs belong to the same PC subtype. More experiments are needed to elucidate whether there is a difference between CTR-PCs and SND-PCs not only in their spatial organization but also with regards to their molecular make-up.

### Functional implications

The recurrent CENTER/SURROUND antagonistic organization of PCs revealed by our experiments appears to be well-suited for a number functions associated with optimal motor control: (1) Single effector dynamic control (Fig. 6a): In this architecture, CTR- and SND-PCs project to the same motoneuron pool via separate populations of CN neurons. Suppression of SS firing in CTR-PCs can drive the effector ‘A’ and increase of firing in SND-PCs can terminate the drive (Chen and Evinger, 2006; Herzfeld et al., 2015). (2) Agonist/antagonist control (Fig. 6b): CTR-PCs control the motoneuron pool for the agonist muscle of effector ‘A’, and SND-PCs control the motoneuron pool of the antagonist muscle. Suppression of firing in CTR-PCs contracts the agonist to accelerate ‘A’ and increase of firing in SND-PCs relaxes the antagonist to support the acceleration (Hore et al., 1991; Sanchez-Campusano et al., 2012; Freeman, 2015; Becker and Person, 2019; De Zeeuw, 2021). (3). Lateral inhibition (Fig. 6c): CTR-PCs and SND-PCs are connected to different effectors, ‘A’ and ‘B’. CTR-PCs drive effector ‘A’ while SND-PCs suppress the movement of effector ‘B’. Examples of this architecture come from parts of the brain such as the basal ganglia, which are thought to control movement by driving a desired specific action while suppressing other undesirable actions (Mink, 1996; Hikosaka et al., 2000; Klaus et al., 2019). (4). Forward model (Fig. 6d): While CTR-PCs drive effector ‘A’, SND-PCs receive the copy of the motor command and predict the sensory consequence of the movement (Wolpert et al., 1998; Pasalar et al., 2006; Laurens et al., 2013; Brooks et al., 2015; Therrien and Bastian, 2015; Kawato et al., 2021). The output of the forward model is broadcasted throughout the brain to allow the motor system to generate accurate motor commands very quickly without having to wait for slow proprioceptive sensory feedback signals.

**Figure 6.**
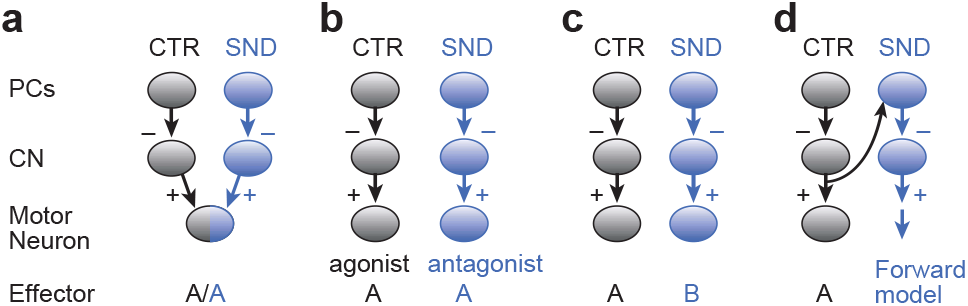
Recurrent CENTER/SURROUND architectures for motor control. **a**, Single effector dynamic control. CTR-PCs project to CTR-CN neurons, SND-PCs project to SND-CN neurons, and both project to a single motoneuron pool responsible for controlling a single effector ‘A’ (corresponding to a particular part of the body). **b**, Agonist/antagonist control. CTR-PCs project to CTR-CN neurons which control motoneurons of the agonist muscle of ‘A’, and SND-PCs project to SND-CN neurons which control motoneurons of the antagonist muscle. **c**, Lateral inhibition. CTR-PCs project to CTR-CN neurons which control motoneurons of effector ‘A’, while SND-PCs project to SND-CN neurons which control motoneurons of effector ‘B’. **d**, Forward model. CTR-PCs project to CTR-CN neurons which control motoneurons of effector ‘A’, while SND-PCs receive the copy of the motor command and project to SND-CN neurons which are the output of a forward model for predicting the sensory consequences of moving effector ‘A’.

In addition to the four functions for motor control listed above, it has been suggested that recurrent circuits between the cerebellum and other parts of the brain could be essential for multiple aspects of motor learning, including: (1) learning preparatory activity during planning (Houk and Wise, 1995; Gao et al., 2018; Li and Mrsic-Flogel, 2020), (2) learning of movement sequences (Hikosaka et al., 1999; Penhune and Steele, 2012; Leggio and Molinari, 2015), (3) learning from sensory-prediction errors when information about motor errors is not available (Jordan and Rumelhart, 1992; Miall et al., 1993; Porrill et al., 2004a; Dean et al., 2010; Ito, 2013), and (4) learning movement dynamics via generation of temporal filters using reservoir computing (Yamazaki and Tanaka, 2007; Rossert et al., 2015). To fully define the role of recurrent signals in the cerebellum during motor control and learning, it will be necessary to selectively manipulate LOOP and SPIRAL circuits between the cerebellum and other brain regions while assessing the impact on the firing of cell-specific neural populations and on behavioral performance.

### Summary and future directions

We investigated the function of the recurrent LOOP and SPIRAL pathways in the cerebellum and made major discoveries that require making revisions to some of the key computational principles underlying the function of the cerebellum. Namely, instead of having cerebellar modules that process information independently of each other along a feedforward pathway, we demonstrate that the function of different cerebellar modules is inexorably tied to each other via recurrent circuits. Our results may shed light on the function of recurrent circuits in many other brain regions, especially those with LOOP and SPIRAL pathways (Hupe et al., 1998; Haber et al., 2000; Nakazawa et al., 2002). In addition, there are few artificial neural networks that contain the type of antagonistic modules with LOOP and SPIRAL connections we have uncovered in the cerebellum. Comparing the capabilities of signal generation and the underlying circuit dynamics in modular architectures with and without LOOP and SPRAL connections could lead to new breakthroughs in machine learning with brain-like artificial neural networks.

## ONLINE METHODS

### Animals

All procedures were approved by the Baylor College of Medicine Institutional Animal Care and Use Committee based on the guidelines of the US National Institutes of Health. Experiments were performed on male C57BL/6J (*n* = 8 mice), PCP2-ChR2 (*n* = 4), nNOS-ChR2 (*n* = 3), and EAAT4-GFP (*n* = 4) mice at least 12 weeks of age housed on a reverse light/dark cycle (8:00 lights-off to 20:00 lights-on); mice were housed in cages of up to four before surgery and were singly housed after surgery. Ambient temperature was maintained at 20–22.2 °C, and ambient humidity was maintained at 30–70% relative humidity. No statistical methods were used to predetermine sample sizes, but our sample sizes were similar to those reported in previous publications (Halverson et al., 2015; Ohmae and Medina, 2015; ten Brinke et al., 2015). Mice were randomly allocated into different groups.

### Surgery

Procedures have been described previously (Heiney et al., 2018). In brief, mice were anesthetized with isoflurane (5% by volume in O2 for induction, 1–2% by volume for maintenance; SurgiVet) and kept on a heating pad to maintain body temperature. Surgeries were carried out under sterile conditions, and mice received preoperative analgesia (0.02 ml of 0.5% bupivacaine and 2% lidocaine, subcutaneously at incision site; 5 mg/kg meloxicam, subcutaneously). Each mouse received a craniotomy over the right anterior cerebellar cortex, and the implantation of a recording chamber (2.5 mm × 2.5 mm; protected by a three-dimensionally printed chamber, NeuroNexus) above the craniotomy and a head-plate, secured to the skull using two screws and C&B Metabond.

### Stimulus control and behavioral procedures

In all experiments, mice were head-fixed on top of a cylindrical treadmill and allowed to walk freely (Heiney et al., 2014b; Heiney et al., 2018). Facial and eyelid movements ipsilateral to the cerebellar region of interest were monitored with a high-speed monochrome camera (GE680, Allied Vision) under infrared illumination. Video frames (200 fps) were triggered by a Sys3 processor (RZ5, TDT) and stored with the MATLAB Video Acquisition Toolbox. A Sys3 processor (RZ5, TDT) was used to control the timing of stimuli during conditioning. The US was an airpuff of nitrogen (20-40 psi, 10–20 ms) delivered via a 23-gauge flat-ended needle placed ∼4 mm in front of the right cornea of the mouse. The CS were either a 500-ms blue light LED positioned 8 cm in front of the mouse’s face, or a 500-ms tone of white noise delivered via a speaker (4-Ω magnetic speaker, FF1, TDT) positioned 20 cm in front of the mouse. The volume of the white noise was set before the first conditioning session to just below the threshold for eliciting short-latency startle movement of the eyelid. The interstimulus interval between the CS and US was 200 ms except one mouse (370 ms). For mice other than PCP2-ChR2 mice, US-omitted trials were used to observe the CR driven by the CS without contamination from the reflex eyeblink response after the US. The fraction of the US-omitted trial was ∼25% of all the conditioning trials, but the fraction was increased in a few sessions to collect data with variation in CR size needed for analysis. The minimum interval between trials (intertrial interval; ITI) was 20 seconds for sessions with photostimulation and 7–16 seconds for the other sessions, but trials could only start if the eyelid position was stable for at least 0.6 s. During sessions with photostimulation, light was delivered using analog-controlled, 473-nm lasers (Blue Sky Research, FTEC2473) and a 200-μm-diameter patch cable (ThourLab M86L01).

### Behavioral analysis

Movement of the eyelid was calculated frame by frame by counting the number of white pixels in a thresholded binary image of the eye and surrounding fur, according to a procedure described previously (Heiney et al., 2014b). Eyelid closure was measured in units of fraction eyelid closure (FEC), ranging from 0 (fully open) to 1 (fully closed). Trials in which FEC did not reach at least 0.1 in the ISI period were defined as no-CR. CR onset was defined as the time when the low-pass filtered eyelid velocity trace reached a threshold of 0.002 FEC/ms. For an Extended Data Figure, UR onset in no-CR trials were defined with the same threshold.

### Single-unit recording

Extracellular recording of SSs and CSpks in Purkinje cells was performed with 1–5 MΩ tungsten microelectrodes (75 μm of shaft diameter, FHC), glass capillary electrodes (BF150-86-10, Sutter instrument) with 2–7-μm tip and 3–6-MΩ impedance (P-1000, Sutter instrument), or a tetrode (Thomas Recording, AN000968). The electrodes were controlled with a hydraulic microdrive (MMO-220A, Narishige) mounted on a three-axis micromanipulator (WPI, M325; Narishige, SMM-100). The voltage signal was acquired at a 24,414-Hz sampling rate, and band-pass filtered between 20–10,000Hz using a digital processor (RZ5, TDT). The electrodes were directed along a 10- or 15-deg angled axis from posterior dorsal to anterior ventral, relative to the vertical plane. In addition, the data of CSpks of Purkinje cells of wildtype mice and the recording data of the deep cerebellar nucleus includes data from previously collected datasets (Ohmae and Medina, 2015; Ten Brinke et al., 2017).

### Electrophysiology analysis

We recorded a total of 138 Purkinje cells (61 from wildtype mice trained with the 220-ms ISI, 17 cells from a wildtype mouse trained with the 370-ms ISI, 40 cells from PCP2-ChR2 mice, 11 cells from nNOS-ChR2 mice, and 9 cells from EAAT4-GFP mice). The spike sorting and the electrophysiological identification of Purkinje cells was performed as described previously (Ohmae and Medina, 2015; Kim et al., 2020).

The spike density of SS was constructed by convolving a Gaussian function with the SS events in each trial. The width of the Gaussian function was 5 ms for the analysis of activity onsets (in Figure 5 and in Supplementary Figure 10) and 10 ms for all other analyses. A CSpk peristimulus time histogram (PSTH) was constructed for each Purkinje cell with 10-ms bins. The spontaneous (baseline) firing rates were defined for SSs and CSpks in the 500-ms time window before the trials started.

CENTER Purkinje cells are defined as those with a CSpk response after the unexpected puff (measured in 120-ms time window after the puff) that is at least 4 times the spontaneous rate. On the other hand, SURROUND Purkinje cells were defined as those with a CSpk response after the unexpected puff that was less than 4 times the spontaneous rate and a SS response after the unexpected puff (measured as the peak of the spike density in the 100-ms time window after the puff and referred to as (ΔSS, puff) in Figure 1h and 3h) that was at least 20 Hz. The remaining recorded Purkinje cells had a Cspk response less than twice the spontaneous rate after the unexpected puff and a SS response less than 20 Hz after the unexpected puff; they exhibited little or no CR-related activity (displayed only in Fig. 3g,h with gray trace and dots) and were excluded from further analysis. For Figure 1f,h and 3f,h, the ΔSS during CR was defined as the trough (for CENTER Purkinje cells) or the peak (for SURROUND Purkinje cell) of the spike density in the CS-US interval. For Figures 2g,h and 4f,g, the spike rates in Hz were measured by averaging spike counts in each time window across trials and dividing by the bin size (0.07 sec).

To predict the spike density of SS with the kinematics of eyelid movement, we used a linear model: SS(t) = K1* position(t-tau) + K2 * velocity(t-tau) + K3 * acceleration (t-tau) + K4. The fitting parameters were optimized to minimize the total squared error for the three population averaged spike density traces in small-CR, middle-CR, and big-CR trials for which the conditioning trials were divided according to the size CR (as in Supplementary figure 3a,d). The fitting duration ranged from -100 ms to 750 ms relative to the CS onset.

During rAIN photoinhibition, the time course of CR-related responses in SS firing of CTR-PCs (Fig. 2f) and SND-PCs (Fig. 4e) was calculated by subtracting the SS response in conditioning trials with laser from the SS response to the laser alone. Supporting this linear operation, the information integration of the PC has been reported to be extremely linear (Walter and Khodakhah, 2009). For SND-PCs, since the antidromic activation was not apparent on average, the presence or absence of subtraction did not affect any of the results.

For detecting the onset of the neural response for each Purkinje cell and CTR-CN neuron relative to the CR onset, we normalized the spike density (5-ms gaussian filtered) such that the baseline (spontaneous) rate is 0 and the peak rate (for SND-PCs and CTR-CN) or the minimum rate (for CTR-PCs) is 1. The last time point between -100 ms and the time at which the normalized response reached 1 with a value less than 0.05 was defined as the onset of activity.

### Photostimulation experiments

For rAIN photoinhibition, we photostimulated the GABAergic axon terminals of Purkinje cells at rAIN, using PCP2-ChR2 (Purkinje cell specific ChR2 under L7 promotor) mice as previously reported (Heiney et al., 2021). The following two methods to suppress rAIN while recording Purkinje cells in parallel were less successful: First, we tried to infuse muscimol or lidocaine into rAIN (pump or iontophoresis), but the infusion site was so close to the Purkinje cells being recorded (about 0.6 mm) that the direct diffusion of muscimol/lidocaine to the Purkinje cells affected their spontaneous firing rates even when the infusion volume was strictly controlled to the minimum required to suppress CR. Second, we attempted Arch expression in rAIN with AAV5-hSyn-Arch but abandoned it because the toxicity of AAV5-hSyn-Arch could interfere with learning (Kim et al., 2020) and most of the rAIN cells died in the pilot experiment.

Prior to all sessions, the site of photostimulation in rAIN was identified using a previously reported method with electrical stimulation (Heiney et al., 2014a; Heiney et al., 2018) and the stereotaxic coordinates were recorded. In each daily experiment, after the recording electrode was inserted into the brain through the dura matter, the optic fiber with the sharpened tip (200-μm core diameter, ThorLabs AFS200/220Y, 0.22 NA) was inserted into the brain according to the recorded coordinates and lowered to just dorsal to rAIN (1800 μm from the surface). Before lowering the recording electrode and finding Purkinje cells, the photostimulation site was evaluated again as a daily assessment, in which the inhibition of CR by the photostimulation was measured. If the suppression of the CR was not sufficient (<90%), the stimulation site was systematically adjusted by trial and error (in 100 μm steps). Sessions in which CRs were not sufficiently suppressed despite this procedure were excluded from further analysis. The angles of the optic fiber and electrode (see Single-unit recording section) were kept at 15° and 10°, respectively, throughout all sessions. This 5-degree difference in angle allowed us to load the electrode and fiber holders (Narishige, UPN-10) on the micromanipulators (WPI, M325; Narishige, SMM -100) without interference.

Four types of trial were interleaved and tested in each photostimulation session: (1) normal eyeblink conditioning trials with CS and US; (2) eyeblink conditioning trials with photo-inhibition which contained CS, US, and laser stimulus (350-ms duration starting 10 ms after tone CS onset or 40 ms after LED CS onset); (3) laser-alone trials (350-ms duration); (4) US-alone trials. Laser intensities ranged from 15 to 40 mW (the most frequent power was 30 mW). Two mice were trained with both tone and LED CSs. One mouse was trained with a tone CS, tested, and then retrained with LED CS and tested again. The other was trained with the LED CS first and then with the tone CS. During the session with tone CS, blue ambient illumination (10×, Uxcell a14122700ux0012 in series, 550-Ω resistance to ground, 0.5 A power supply by Tekpower TP3005N) was present throughout the sessions to mask light escaping from the junction between the patch cable and implanted ferrule during photostimulation. For the LED CS, we pulled out the patch cable and confirmed that any light leak had no effect on the CR.

### Histology

The location of a few CENTER (n = 5) and SURROUND (n = 7) Purkinje cells was marked with electrical microlesion (0.01 mA, 20 sec) through Thomas tetrode or with dye injection (2% Pontamine sky blue, Sigma 2610-05-1; 0.2% Alcian blue) through glass pipette by picosplitzer (World precision instruments, pv820, 20-40 PSI, 20 ms). The brain was extracted and sectioned with 40-micrometer thickness 7-10 days after the marking of the microlesion and immediately after the marking of dye injection. The marking was detected by red fluorescent Nissl (3.3% in PBS, ThermoFisher, N21482) or immunohistochemistry (anti-mouse secondary antibody) for the electrical microlesion, or by direct observation without staining for the injected dye. The zebrin zones were visualized using immunohistochemistry with zebrin-antibody (gift from Dr. Richard Hawkes and Dr. Roy V. Sillitoe) for wildtype mice, or using a fluorescent microscope (Zeiss, Axio-Imager microscope and ZEN software, blue edition) for EAAT4-GFP mice. After mice completed the experiment of photo-inhibition of CTR-DCN (rAIN), the location of the optic fiber tip was marked with intense light stimulation through the optic fiber (Shanghai Laser & Optics Century, BL473T3-150FC, 85 mW, 45 sec for two mice and 120 sec for two mice). The brain tissue was extracted 3 days after the marking and sectioned. The marking was detected by cresyl violet (0.5% cresyl violet and 0.3% glacial acetic acid in distilled water) and the bright-field mode of the microscope.

### Statistics and reproducibility

All statistical analyses were performed in MATLAB (v.2015a, including Statistics and Machine Learning Toolbox). We used nonparametric statistical tests without assuming normality, except for data in Figure 1f,h and 3f,h, where we used Pearson’s correlation coefficient. All tests were two-sided. No randomization was used, but mice were assigned to specific experimental group without bias and no animals were excluded. The experimenters were blind to task condition and behavior during spike sorting. Analyses were based on automated scripts applied across experimental conditions and thus were not subject to any experimenter bias. All electrophysiological and behavioral experiments with wildtype and PCP2-ChR2 mice were conducted over two rounds, with each experiment conducted on a different group of mice and producing the same results (Figs. 1–5 and Supplementary Figs. 1, 3–6, 8-9).

## KEY RESOURCES TABLE

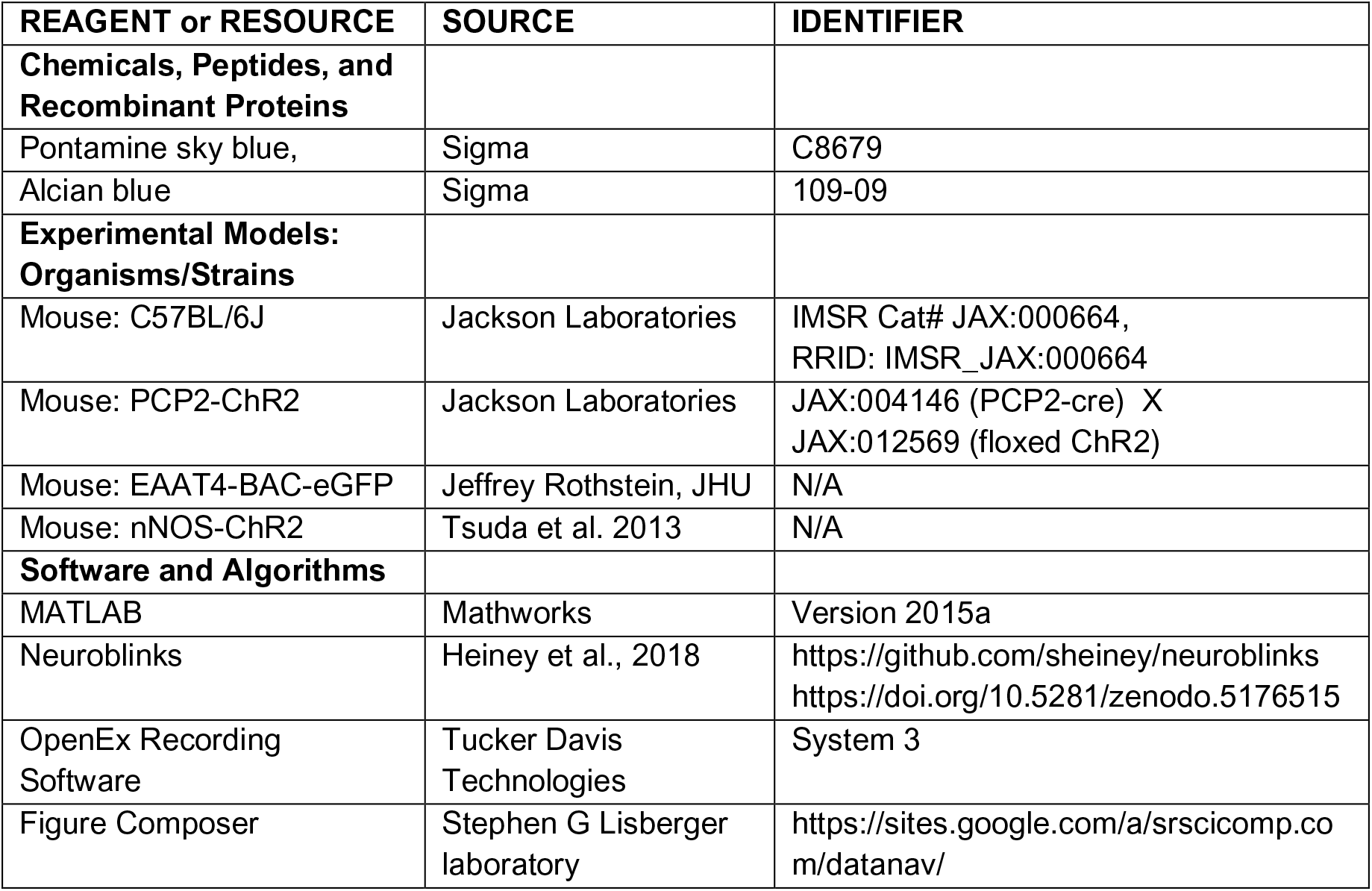

## RESOURCE AVAILABILITY

### Lead contact

Further information and requests for resources should be directed to and will be fulfilled by the Lead Contact, Javier Medina.

### Materials availability

This study did not generate new unique reagents.

### Data and code availability

- The custom software for controlling the task and analyzing the eyeblink behavior is available on Github (https://github.com/sheiney/neuroblinks).
- Any additional information required to reanalyze the data reported in this paper is available from the Lead Contact upon request.

## ACKNOWLEDGMENTS

We thank Dr. Jeannie Chin for help with immunohistochemical procedures, Dr. Richard Hawkes and Dr. Roy V. Sillitoe for a generous gift of anti-zebrin antibody, Dr. Jeffrey Rothstein for a generous gift of EAAT4-BAC-eGFP mice, and Dr. Tadashi Kodama and Dr. Hirofumi Fujita for instructions on the dye injection procedure. Supported by grants to JFM from the National Institutes of Health (R01 MH093727; RF1 MH114269; NIH R01NS112917) and to SO from the National Institutes of Health (R34 NS118445).

## AUTHOR CONTRIBUTIONS

Conceptualization, S.O. and J.F.M; Data curation, S.O. and K.O.; Formal Analysis, S.O.; Funding Acquisition, J.F.M and S.O.; Investigation, S.O., K.O., S.A.H., and D.S.; Methodology, S.O. and J.F.M. Project administration, S.O. and J.F.M.; Resources, J.F.M and S.O.; Software, S.O.; Supervision, J.F.M.; Validation, S.O., K.O., and S.A.H.; Visualization, S.O., J.F.M., and K.O.; Writing – original draft, S.O. and J.F.M.; Writing – review & editing, J.F.M. and S.O.

---

**Supplementary figure 1, related to Figure 1.**
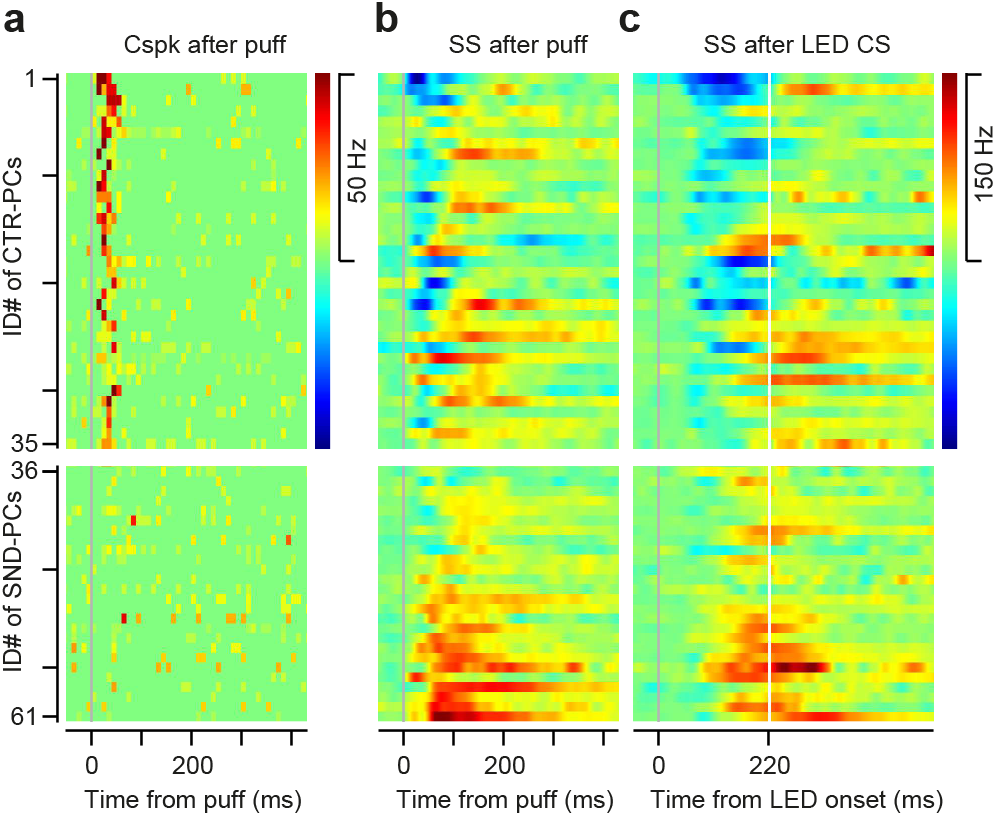
Summary of eyepuff-driven and CR-related responses of individual Purkinje cells. **a,** The CSpk responses to the unexpected puff of individual CENTER (top, n = 35) and SURROUND (bottom, n = 26) Purkinje cells (histogram with 10-ms bins). **b**, The SS responses to the unexpected US (spike density with 10-ms Gaussian). **c**, The CR-related activities of SS. The spike density after the time of the US (white vertical line) is computed by analyzing trials in which the US was omitted.

**Supplementary figure 2, related to Figure 1.**
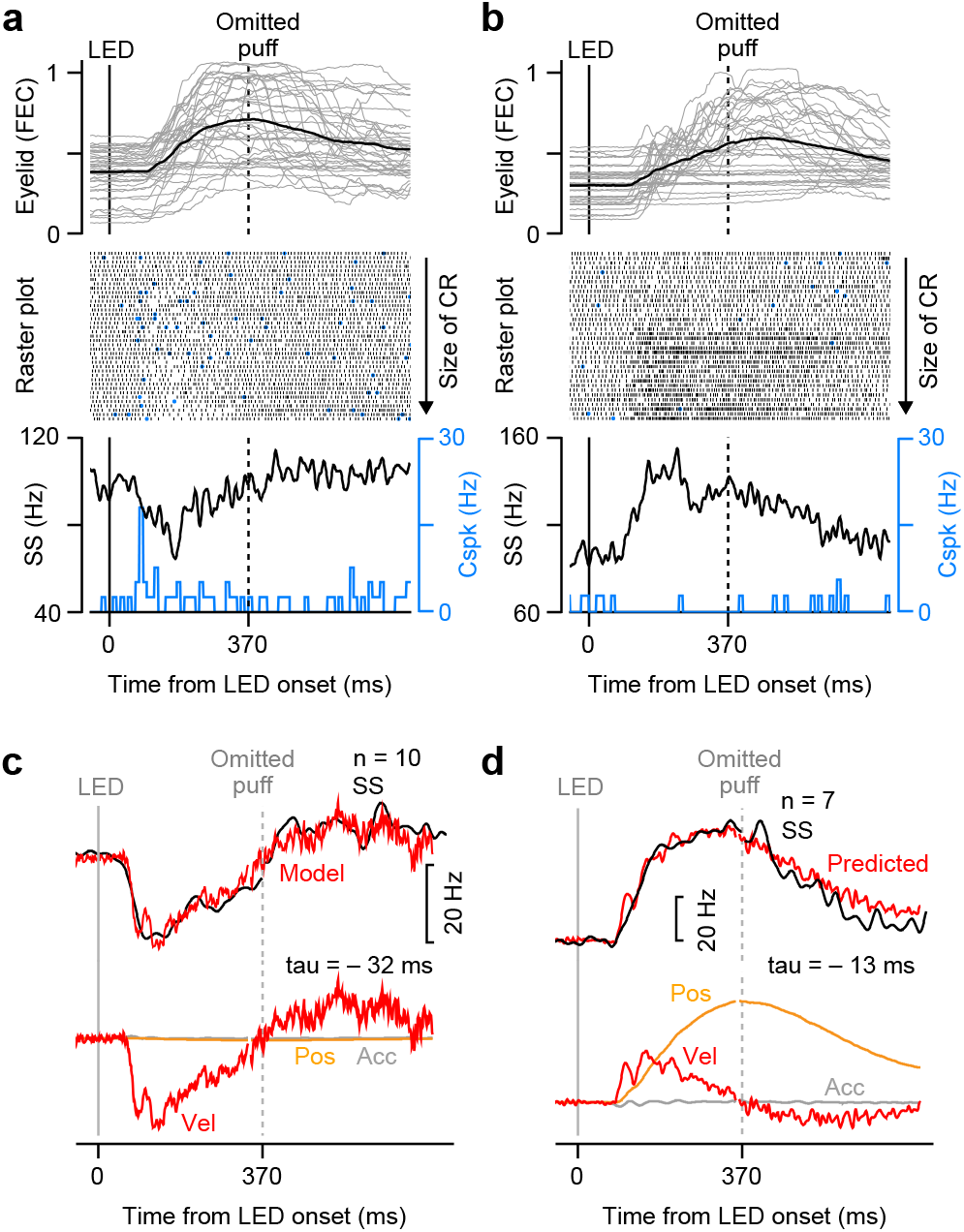
CR-related activities of Purkinje cells in mice trained with a longer CS-US interval. **a-b,** The Eyelid movements, and SSs (black) and CSpks (blue) in US-omitted trials of representative CENTER (**a**) and SURROUND (**b**) Purkinje cells, trained with a long CS-US interval (370 ms). **c-d**, Top, Model prediction for population averaged SS firing of CENTER (**c**) and SURROUND (**d**) Purkinje cells with the kinematics of the eyelid movement: SS(t) = K1* position(t-tau) + K2 * velocity(t-tau) + K3 * acceleration (t-tau) + K4. Bottom, Contribution of each kinematic component to the predictive model.

**Supplementary figure 3, related to Figure 1.**
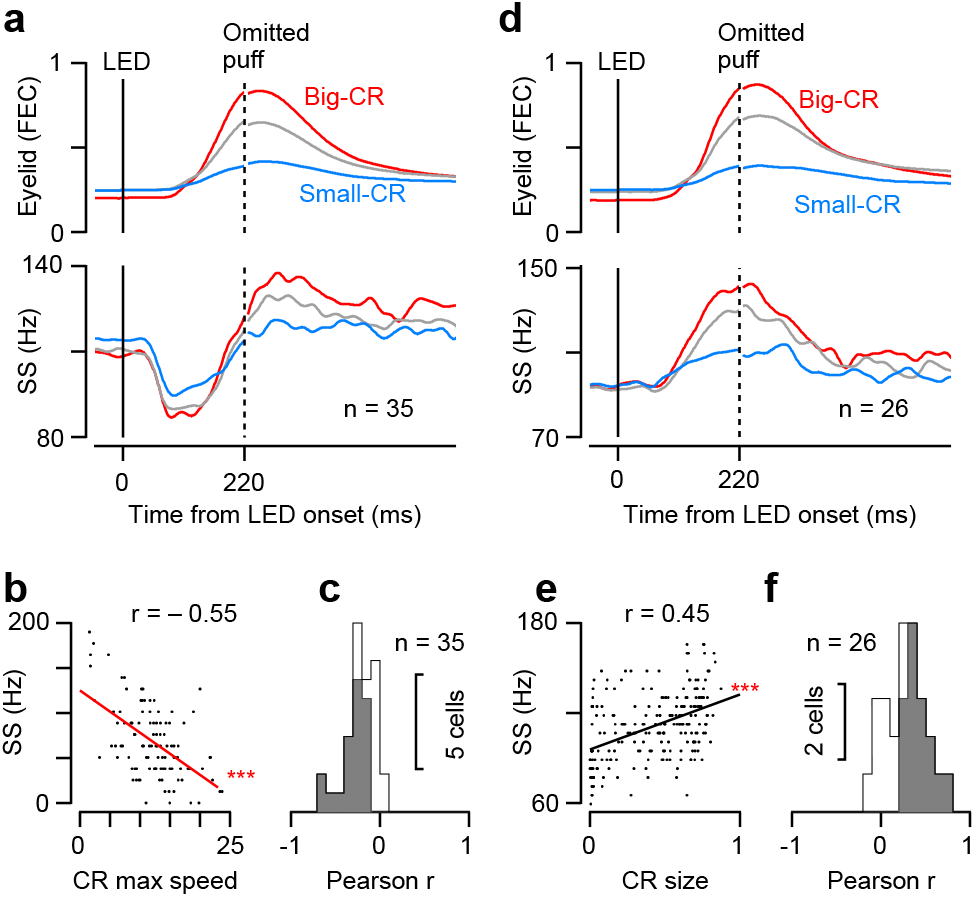
Relationship between CR-related SS response of Purkinje cells and CR kinematics. **a,d,** The size and time course of population-averaged CR-related activity of CENTER (**a**) and SURROUND (**d**) Purkinje cells in trials with CRs of different sizes. **b,** The trial-by-trial correlation between the maximum speed of the CR and the SS firing rate in a 80-ms time window of the representative cell in Figure 1 (Pearson’s correlation coefficient, r = -0.55, P < 0.001). The time window was adjusted to the time where SS rate of the cell was minimum (80-ms interval starting 100 ms after the CS onset). **e**, The trial-by-trial correlation between the size of the CR and the SS firing rate in a 150-ms time window (starting 50 ms after the CS onset) of the representative cell in Figure 3 (Pearson’s correlation coefficient, r = 0.45, P < 0.001). **c,f**, Pearson’s trial-by-trial correlation computed as in **b** and **e** for all CENTER (**c**) and SURROUND (**f**) Purkinje cells. The cells with significant correlation are displayed with gray shade (**c**, n = 21/35 cells; **f**, n = 15/26 cells).

**Supplementary figure 4, related to Figure 2.**
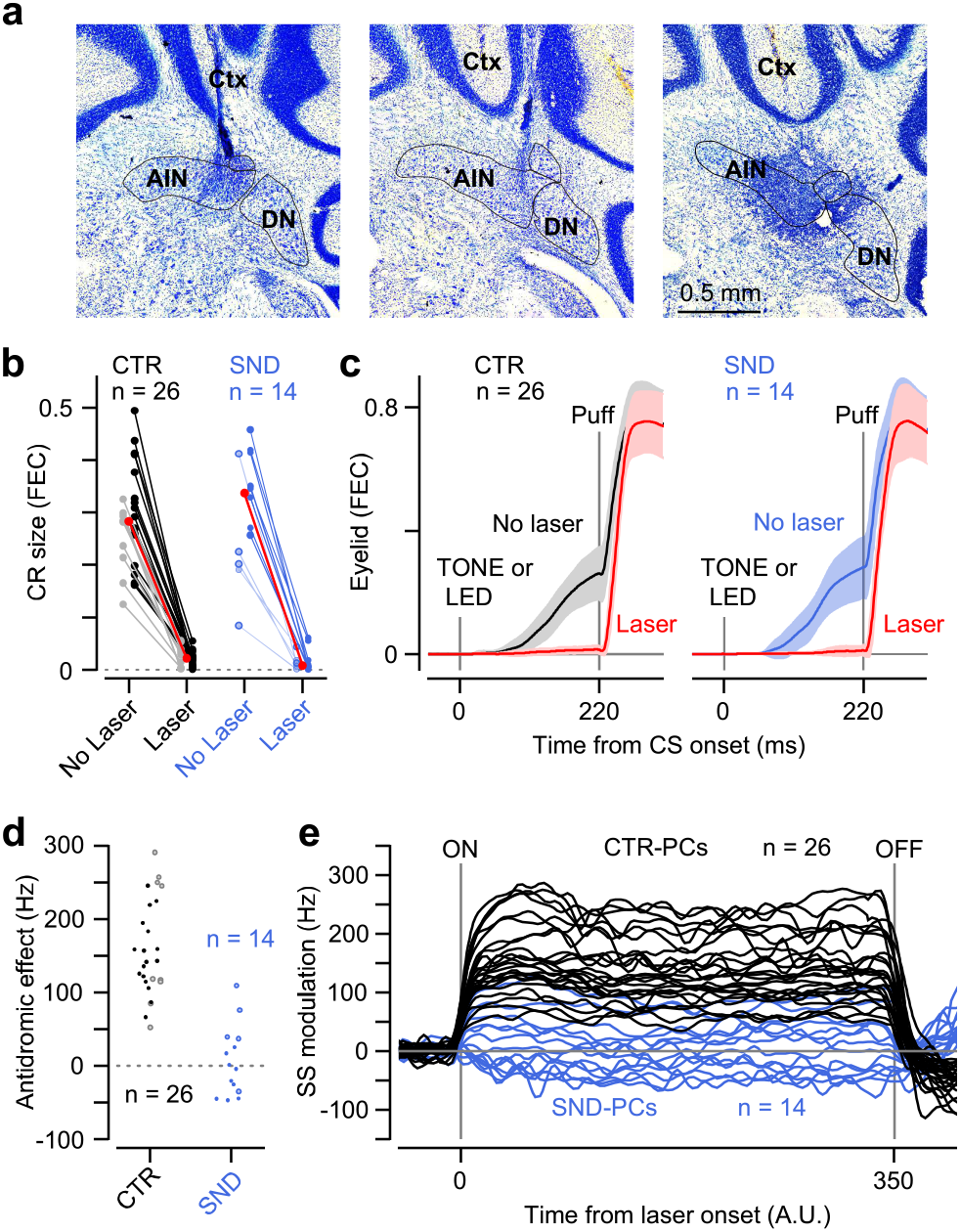
Validation of method for inhibition of CENTER module output neurons in rAIN. **a,** Locations of fiber tips, identified with marking microlesions created by high-intensity laser stimulation in individual mice (Nissl staining; n = 3/4 mice) AIN, anterior interpositus nucleus; DN, dentate nucleus; Ctx, cerebellar cortex. **b**, Size of CRs in the absence and presence of photostimulation of GABAergic axon terminals in rAIN (CENTER CN) for individual sessions in which CENTER (black) and SURROUND (blue) Purkinje cells were recorded (darker colors for LED CS, brighter colors for TONE CS, red for the median). **c**, Eyelid movements (mean ± s.d.) in the absence or presence of the photostimulation. **d**, Antidromic-driven SS response during photostimulation for individual CENTER and SURROUND Purkinje cells. **e**, SS activity of CENTER and SURROUND Purkinje cells during photostimulation. In the majority of cells, the duration of photostimulation was 350 ms (n = 22/26 for CTR-PCs, n = 10/14 for SND-PCs).

**Supplementary figure 5, related to Figure 2.**
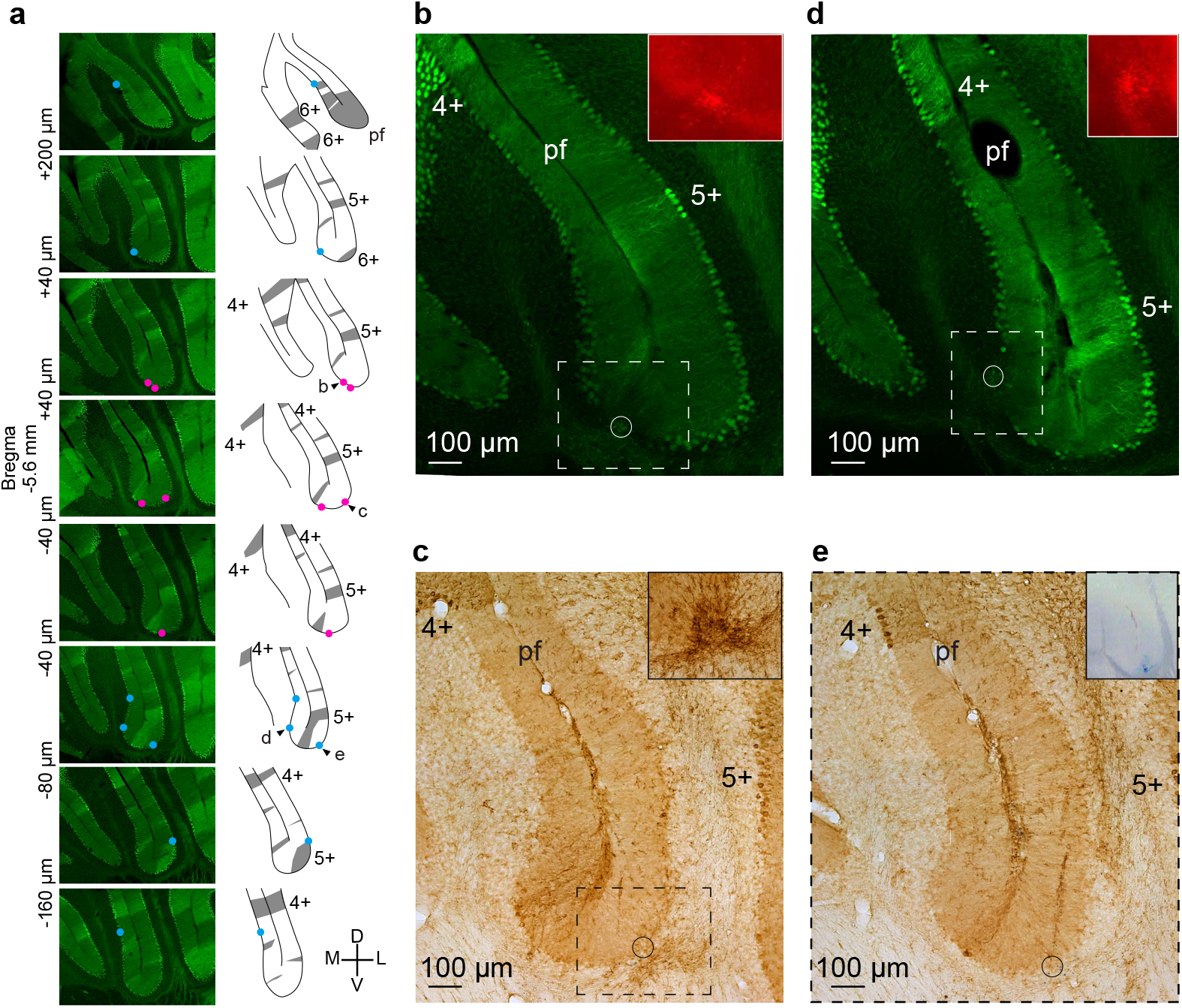
Locations of CENTER and SURROUND Purkinje cells relative to zebrin stripes. **a**, The location of CENTER (red dots; n=5) and SURROUND cells (blue dots; n=7) are shown on coronal sections near the right primary fissure (pf) of a template brain from EAAT4-GFP mice expressing GFP in zebrin-positive Purkinje cells. The sections are ordered from rostral to caudal. Schematic diagram shows zebrin positive regions (gray). The fourth, fifth, and sixth zebrin bands are labeled 4+, 5+, and 6+. **b-e**, Locations (circles) of 2 CENTER (**b,c**) and 2 SURROUND (**d,e**) cells located in zebrin negative regions, visualized by GFP (**b,d**) or immunohistochemistry (**c,e**). Locations were marked with electrical microlesion and detected by fluorescent Nissl (insets, **b,d**) or immunohistochemistry (**c**). The location of the cell in (**e**) was marked with injected dye. Each inset corresponds to the area in dotted square (**b-d**) or the whole field (**e**).

**Supplementary figure 6, related to Figure 2.**
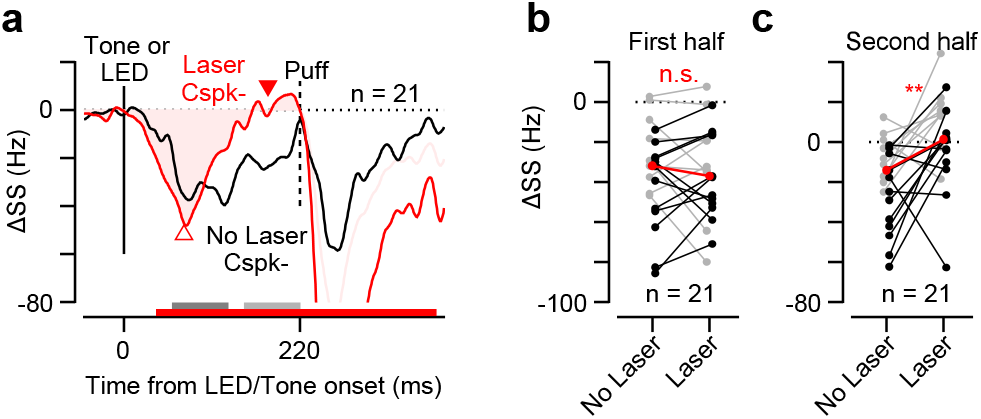
Effect of rAIN photo-inhibition on CENTER Purkinje cells in trials without a CS-driven CSpk. To test whether the effects described in Fig. 2 depend on the presence of CS-driven CSpk’s (Ohmae and Medina, 2015), the same analysis was repeated after excluding trials in which CSpk was present during the 70-ms time period starting 50 ms after LED onset or 50-ms time window starting 20 ms after TONE onset. **a**, Population-averaged CR-related activity of SS in the absence (black) and presence (red) of rAIN photoinhibition. **b,c**, Comparison of firing modulation for individual Purkinje cells and the median (red) after the LED CS (black) and TONE CS (gray) during the first half (**b**, dark gray in **a**) and the second half (**c**, light gray in **a**) of the CR period (Wilcoxon signed rank test; **b**, n = 21 cells, P = 0.69; **c**, n = 21 cells, P =0.0057). n.s. not significant, ** P < 0.01.

**Supplementary figure 7, related to Figure 3.**
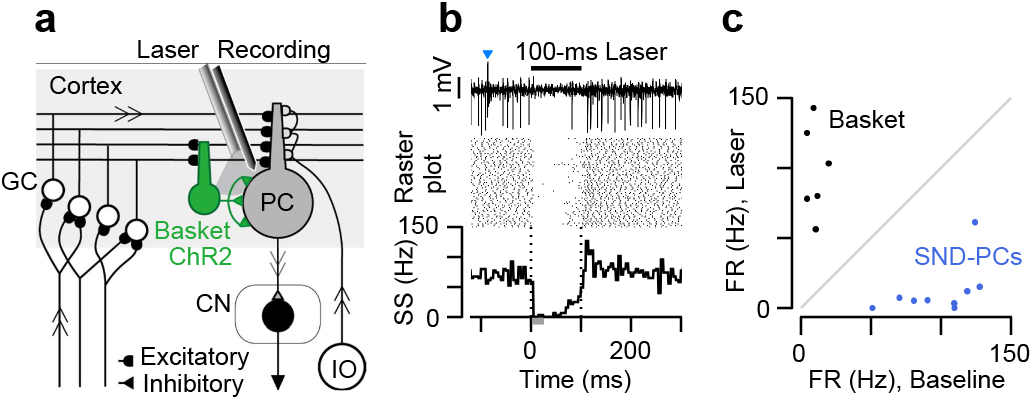
Optogenetic confirmation of Purkinje cells. **a**, Schematic diagram of the method for identifying Purkinje cells using molecular-layer-interneuron specific ChR2 mice (nNOS-ChR2 BAC transgenic mouse line). CN, Cerebellar nuclei; GC, Granule cell; IO, Inferior Olive; PC, Purkinje cell. **b**, Raw extracellularly recorded voltage signal and activity of a representative cell that was identified as a Purkinje cell based on the presence of spontaneous CSpk’s (blue arrow), as well as the suppression of activity during the photostimulation of nearby molecular layer interneurons. **c**, Comparison of firing rates of individual SURROUND Purkinje cells (blue dots) and putative basket cells (black dots) before and during the photo-stimulation (measured in 20-ms time window 5 ms after laser onset, gray bar in **b**).

**Supplementary figure 8, related to Figure 3.**
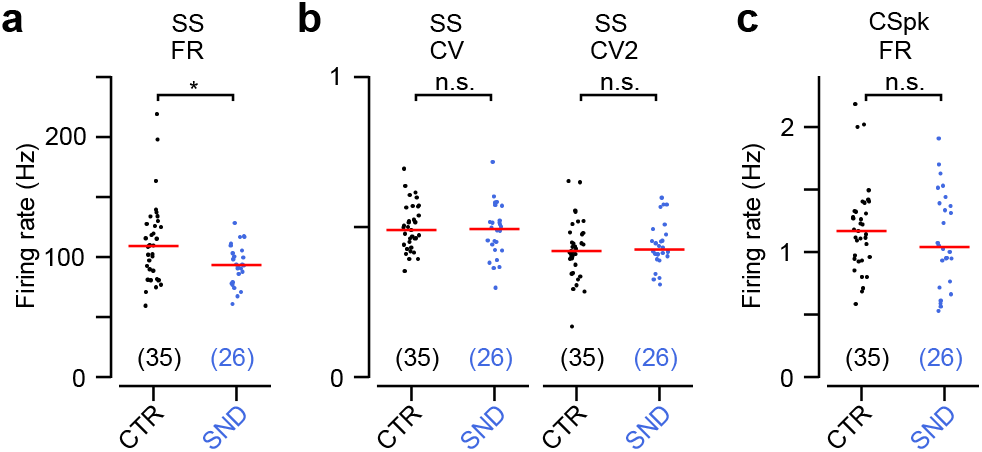
Spontaneous firing properties of CENTER and SURROUND Purkinje cells. **a,** Spontaneous SS firing rates of individual CENTER and SURROUND Purkinje cells (Wilcoxon rank sum test, n = 35 cells and n = 26 cells, P = 0.030). **b**, Regularity of spontaneous SS firing of individual CENTER and SURROUND Purkinje cells, measured by coefficient of variation (CV, Wilcoxon rank sum test, n = 35 cells and n = 26 cells, P = 0.68) and average coefficient of variation for consecutive interspike intervals (CV2, Wilcoxon rank sum test, n = 35 cells and n = 26 cells, P = 0.58). **c**, Spontaneous CSpk firing rates of individual CENTER and SURROUND Purkinje cells (Wilcoxon rank sum test, n = 35 cells and n = 26 cells, P = 0.49).

**Supplementary figure 9, related to Figure 4.**
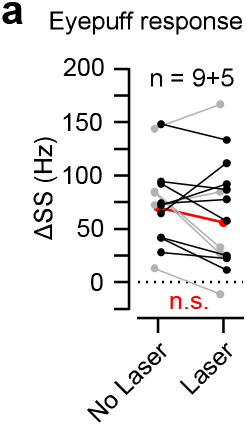
SS response of SURROUND Purkinje cells to the eyepuff during rAIN photoinhibition. **a**, Firing modulation for individual SURROUND Purkinje cells in mice trained with an LED CS (black) or TONE CS (gray) and the median modulation (red) during a period after the eyepuff US (70-ms window starting 50 ms after US onset; Wilcoxon signed rank test, n = 14 cells, P = 0.22).

**Supplementary figure 10, related to Figure 5.**
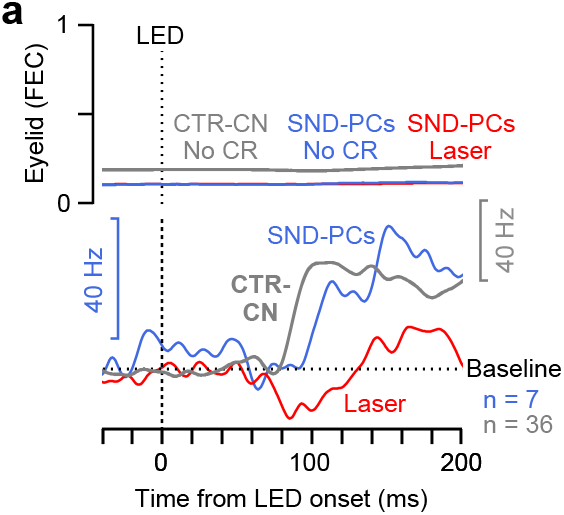
CTR-CN neurons are necessary for the responses of SND-PCs in trials without a CR. **a,** The activity of SND-PCs (blue) and CTR-CN neurons (gray) in trials in which the mouse failed to make a CR (no-CR trials), and the activity of the same SND-PCs in trials with photoinhibition of CTR-CN neurons in rAIN (red). Note that despite the absence of motor output in no-CR trials, there was a sequential activation of CTR-CN neurons first followed by SND-PCs, and that the firing rate increase of SND-PCs was largely suppressed by the photoinhibition of CTR-CN, suggesting that the response of SND-PCs is driven by CTR-CN and does not require motor output. The analyzed populations were 9 cells for SND-PCs (from PCP2 mice with LED CS) and 60 cells for CTR-CN neurons. 2 cells in the SND-PCs and 24 cells in CTR-CN neurons were excluded from the analysis because those did not contain enough number of no-CR trials (at least 4 trials).

